# Bursts boost nonlinear encoding in electroreceptor afferents

**DOI:** 10.1101/2024.06.07.597907

**Authors:** Alexandra Barayeu, Maria Schlungbaum, Benjamin Lindner, Jan Grewe, Jan Benda

## Abstract

Nonlinear mechanisms are at the heart of neuronal information processing, for example to fire an action potential, the membrane voltage must exceed a threshold nonlinearity. Even though, linear encoding schemes are commonly used and often successfully describe large parts of sensory encoding nonlinear mechanisms such as thresholds and saturations are well known to be crucial to encode behaviorally relevant features in the stimulus space not captured by linear methods. Here we analyze the role of bursts in p-type electroreceptor afferents (P-units) in the weakly electric fish *Apteronotus leptorhynchus*. It is long known that subpopulations of these cells fire bursts of action potentials while others do not. Previous research suggests, that the non-bursting cells are better at encoding the stimulus time-course while bursting neurons are better suited to encode special features in the stimulus. We here show, based on the analysis of experimental data and modeling, that bursts affect the linear as well as the nonlinear encoding. Theoretical work predicts that in simple leaky-integrate-and-fire model neurons, two periodic stimuli interact nonlinearly when the sum of the two frequencies matches the neuron’s baseline firing rate as quantified by the second-order susceptibility. Indeed, such nonlinear responses have been found in non-bursting P-units when stimulated by two beats simultaneously but only in those cells, that exhibit very low levels of intrinsic noise. In this study, we found that bursts strongly enhance these nonlinear responses which may play a critical role in the detection of weak intruder signals in the presence of a strong female signal, i.e. an electrosensory cocktail party.

## Introduction

Linear approaches are widely and successfully applied to characterize the neuronal encoding performance in many systems (Borst and Haag, 2001). Neuronal information processing, however, is inherently nonlinear; for example whenever the membrane voltage exceeds a threshold nonlinearity, an action potential is fired (Hodgkin and Huxley, 1952; Koch et al., 1995). Nonlinear mechanisms such as thresholds or saturations are well known to be crucial for encoding important features in the stimulus space (Gabbiani et al., 1996) which are not sufficiently captured by linear methods. Here, we are interested in nonlinear encoding. The nonlinearity of a system can be described by calculating higher-order Wiener Kernels e.g. resulting in the second-order susceptibility of the system (French and Butz, 1973; Marmarelis et al., 1999; Neiman and Russell, 2011). From theoretical studies, we know that leaky integrate-and-fire (LIF) neurons show second-order susceptibilities in the weak stimulus regime when stimulated by two periodic stimuli and the sum of the two input frequencies *f*_1_ + *f*_2_ equals *f*_Base_, i.e. the baseline firing rate of the neuron (Voronenko and Lindner, 2017)( Fig. 1). In a previous study, we showed that such nonlinearity can indeed be found in subsets of p-type electroreceptor afferents as well as ampullary electroreceptor afferents of the weakly electric fish *Apteronotus leptorhynchus* (Barayeu et al., 2024). Among the p-type afferents, it was only a subpopulation of very regularly firing neurons that showed elevated levels of nonlinear susceptibility under the same very specific circumstances, when *f*_1_ + *f*_2_ equals *f*_Base_ (yellow bands in Fig. 1) as predicted by theory (Voronenko and Lindner, 2017; Barayeu et al., 2024). These findings make the electrosensory system ideal for testing theoretical predictions of nonlinear stimulus encoding in the weak stimulus regime within the encoding of behaviorally relevant stimuli.

**Figure 1:**
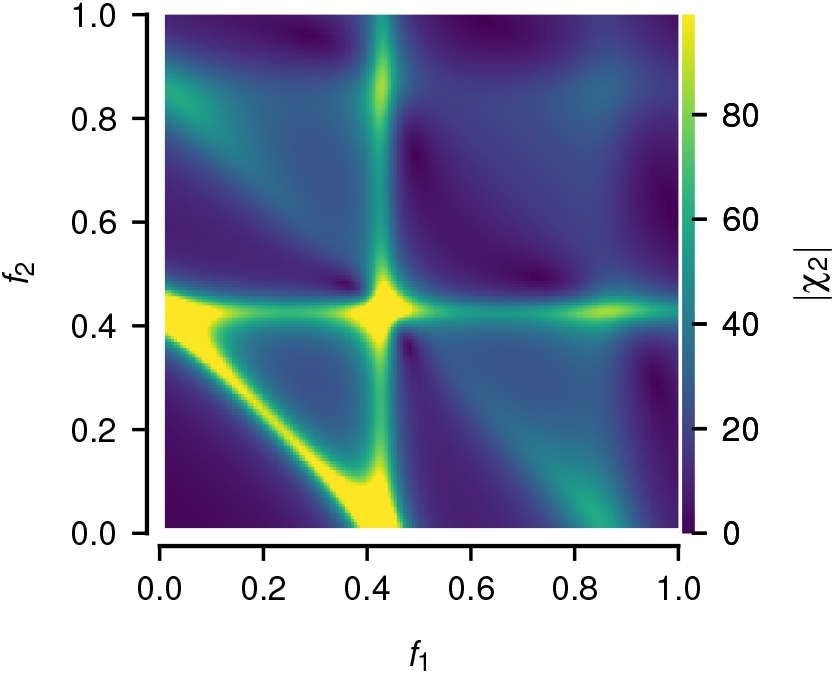
Second-order susceptibility based on the analytic results in Voronenko and Lindner (2017).

These animals are equipped with an electric organ (Salazar et al., 2013), that is constantly active and produces a quasi sinusoidal electric organ discharge (EOD), with a long-term stable and fish-specific frequency (*f*_EOD_, Knudsen, 1975; Henninger et al., 2020). They use their EOD for electrolocation (Fotowat et al., 2013; Nelson and MacIver, 1999) and communication (Zupanc et al., 2006; Fotowat et al., 2013; Walz et al., 2014; Henninger et al., 2018). When two fish are nearby, the EODs interfere and each animal experiences its own field amplitude modulated by the presence of the other. This periodic amplitude modulation (AM) is also called beat and is encoded by the population of p-type electroreceptor afferents (P-units, Bastian, 1981). In interactions of three animals, the animals are exposed to two beats at the same time (Middleton et al., 2006; Stamper et al., 2012; Savard et al., 2011), a situation similar to the above-mentioned stimulation with two periodic signals, for which nonlinear effects are expected (Voronenko and Lindner, 2017). Nonlinear interactions of the two input frequencies may thus be also behaviorally relevant for example to enhance the detection of a weak intruder signal in the presence of a strong foreground signal as observed in freely behaving animals (Henninger et al., 2017). Nonlinear interactions between the stimuli may be the reason why we observed enhanced weak signal detection by the presence of a strong signal in simulations (Schlungbaum and Lindner, 2023). The nonlinear susceptibility was only observed in a small subset of P-units (Barayeu et al., 2024) which is probably too small to have a behaviorally relevant impact. Still, other nonlinear mechanisms might contribute.

One famous nonlinear mechanism is bursting, i.e. the firing of barrages of action potentials that are separated by periods of quiescence. Bursting is observed in many sensory modalities (e.g. Krahe and Gabbiani, 2004; Nowak et al., 2003; Cunningham et al., 2004; Womack and Khodakhah, 2004; Destexhe et al., 1993, just to name a few). The dynamics leading to bursting have been addressed with bifurcation analysis and are well understood (Izhikevich, 2000). Though bursting is also a prominent and long-known feature of subpopulations of p-type electroreceptor afferents (Bastian, 1981; Metzen et al., 2016), their functional role, however, is not so clear. Previous modeling studies concluded that bursting might be a way to establish an additional information pathway. That is, non-bursting P-units are better at encoding the time course of the sensory stimulus while bursting cells are better suited to encode special features in the stimulus (Chacron et al., 2001b, 2004). Interestingly, these cells burst even during their spontaneous activity, i.e. when there are no features to encode. This led to doubts with respect to their functional role. We here analyze the effect of bursting on linear and nonlinear stimulus encoding in P-units of two types of weakly electric fish, *Apteronotus leptorhynchus* and *Eigenmannia virescens*. We observe that bursts enhance the nonlinear susceptibility and widen the range of stimuli that lead to nonlinear interactions. Bursting in conjunction with the known heterogeneity among P-unit properties (Hladnik and Grewe, 2023) may indeed be critical ingredients to explain the weak signal detection in the aforementioned electrosensory cocktail party (Henninger et al., 2018) observed during courtship interactions in freely behaving animals.

## Results

In our previous study (Barayeu et al., 2024) we have worked out the circumstances under which p-type electroreceptor afferents of *Apteronotus leptorhynchus* show nonlinear encoding in the weak stimulus regime. Whether or not a given cell will exhibit an elevated second-order susceptibility can be well predicted by the basic cellular response properties. The CV_Base_ of the interspike interval (ISI) distribution (e.g. Fig. 2 A_ii_ B_ii_) is related to the level of intrinsic noise in the cells and a good predictor for nonlinear encoding in the weak stimulus regime. Cells with a low CV_Base_ will show a |*χ*_2_| matrix with a pronounced peak structure (Fig. 3 A, C). P-units with a high CV do not show such a peakedness of the |*χ*_2_| matrix (Fig. 3 B, C) suggesting that higher CVs lead to less nonlinear coding in the weak stimulus regime.

**Figure 2:**
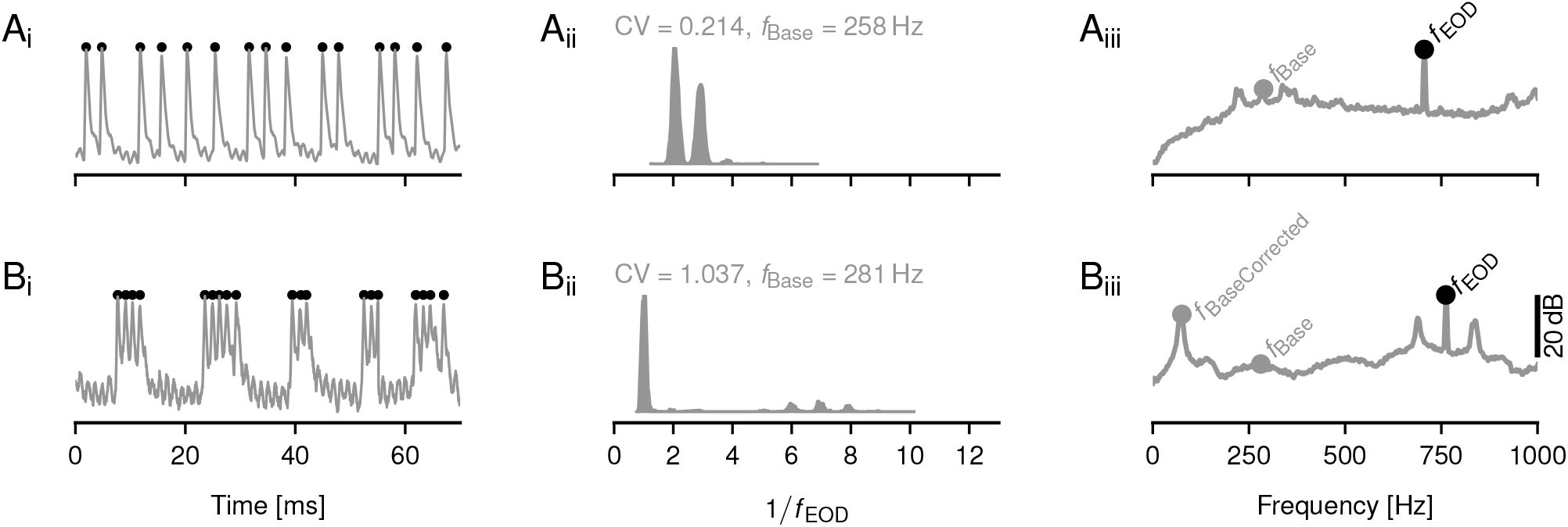
Comparison of the baseline activity of a non-bursty P-unit (A) and bursty P-unit (B). **A**_i_ Voltage trace of baseline activity. The cell is driven only by the self-generated EOD. **A**_ii_ Interspike interval (ISI) distribution of the baseline activity. P-units lock to their own EOD which leads to the characteristic multi-modal distribution of ISIs. **A**_iii_ Power spectrum of the baseline spike trains. Black dot highlights the EOD frequency, the gray dot the baseline frequency of the the recorded P-unit. **B**_i*−*iii_ Same as in A but for a bursty P-unit. In bursty cells action potentials occur in barrages (bursts).

**Figure 3:**
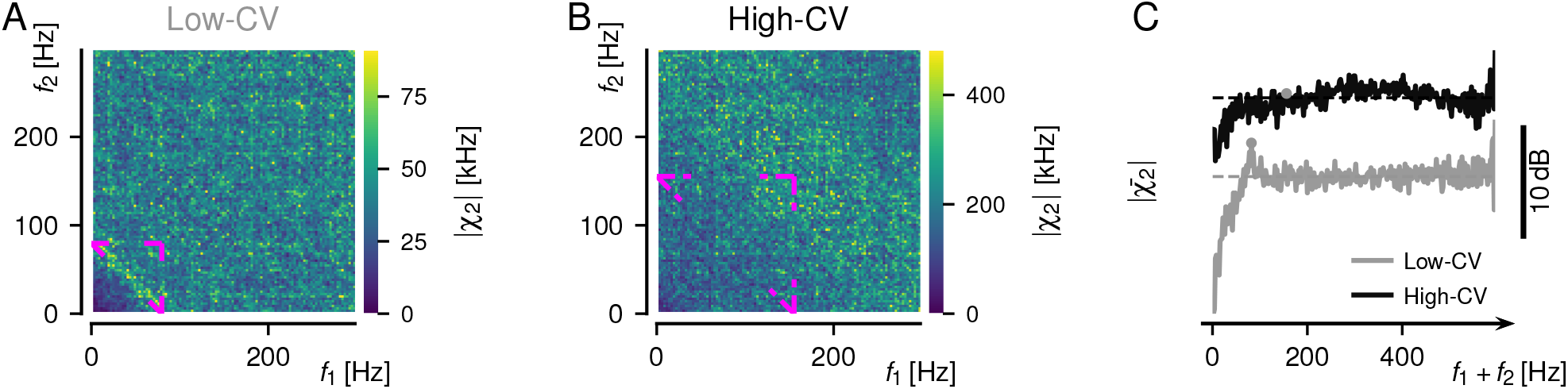
Second-order susceptibility matrices characterizing the nonlinearity of stimulus encoding. **A** |*χ*_2_| matrix of a low-CV P-unit. The second-order susceptibility was estimated at the sum of the frequencies *f*_1_ and *f*_2_ by relating the second-order cross spectrum (8) and the auto-spectra of the stimulus at the given frequencies (Eq. (10)). The matrix was estimated from the neuronal responses to band-limited white noise stimulation. **B** Same but for a high-CV P-unit. **C** Projection of the anti-diagonals onto the main diagonal to characterize the nonlinear structure in the second-order susceptibility matrices. Gray line refers to the low-CV P-unit in A, black to the high-CV P-unit shown in B.

Bursting in the neuronal response leads to very high coefficients of variation of the ISI. From our previous results, one could expect that bursting P-units show less nonlinear encoding (Barayeu et al., 2024). To investigate the role of bursting in nonlinear encoding we compare the cell’s original responses including bursts with a burst-corrected version of the same responses in which only the first spike of each barrage of action potentials was considered (Fig. 4 A). Spikes that occur in a burst of action potentials were identified by the duration of the ISIs (see methods for details). Based on the burst-corrected baseline response the mean burst-corrected firing rate *f*_BaseCorrected_ and the respective CV of the new ISI distribution (CV_BaseCorrected_) were calculated. Burst correction leads to a strongly reduced firing rate and CV_Base_. Interestingly, the power spectrum of the original baseline response shows a distinct peak at *f*_BaseCorrected_ (Fig. 2 B_iii_) but not at the original *f*_Base_. In the stimulus-driven case, the reduced firing rate is reflected in the reduced amplitude of the transfer function used to characterize the linear encoding (Fig. 4 B). The position of the peak and the upper cutoff of the transfer function, however, are similar despite the burst-spike removal.

**Figure 4:**
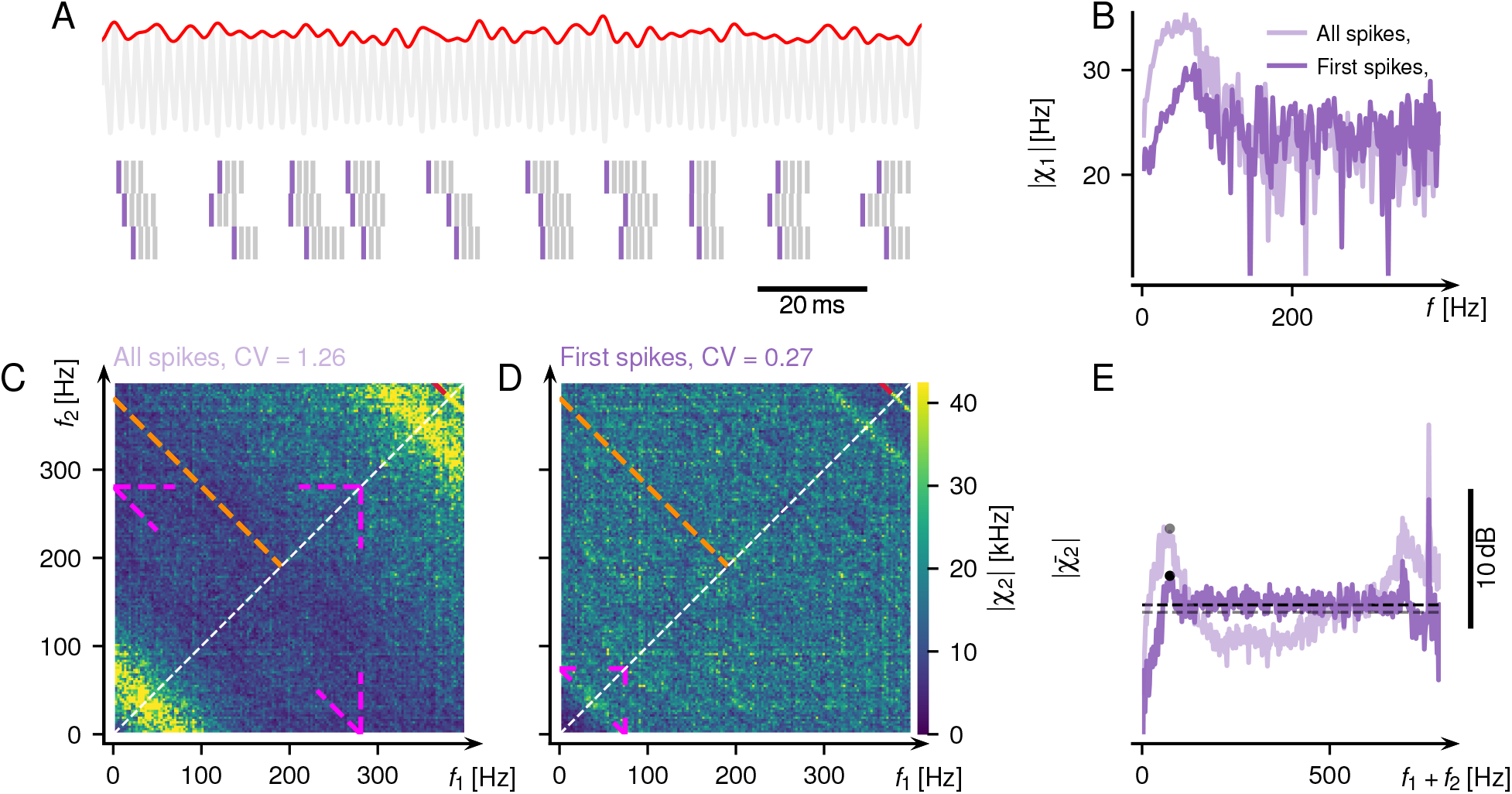
Response of an experimentally measured bursty P-unit before and after burst correction. **A** The cell is driven by the amplitude modulation (red) or the fish’s self-generated electric field which serves as a carrier signal that has an individual frequency for each fish (electric organ discharge, EOD, gray). Spike rasters below show the cell’s bursting response to the stimulus sequence shown above. The first spike of a burst is highlighted in dark. **B** Transfer function (first-order susceptibility) of the cell to the stimulus. The transfer function shows the typical bandpass characteristic, the low-frequency cutoff is a consequence of firing rate adaptation in P-units (Benda et al., 2005). Light color refers to the original (all-spikes) response while dark color shows the transfer function after burst-correction. **C** Second-order susceptibility matrix at *f*_1_ + *f*_2_ (see methods). Bright colors refer to elevated nonlinear response components. Pink lines depict the cases in which theory on LIF models predicts nonlinear responses (Voronenko and Lindner, 2017), i.e. when *f*_1_ + *f*_2_ matches *f*_Base_ (one of the stimulus frequencies is zero in the vertical and horizontal cases). Orange line indicates the Nyquist frequency of the EOD carrier. **D** Same as C but after burst-spikes were removed from the responses. **E** Projection of the anti-diagonals of panels C, D onto the main diagonal to characterize the structure of the |*χ*_2_| matrix. Dots highlight *f*_BaseCorrected_ on the respective curves, dashed lines are the medians. The relative difference between these peak and median |*χ*_2_| values are used to calculate the peakedness of the non-linearity (PNL(*f*_Base_), see methods).

### Bursting boosts second-order susceptibility

In contrast to the expectation that increased CVs would lead to less structure in the second-order susceptibility, the opposite is the case: The second-order susceptibility of the bursty response shows a very pronounced |*χ*_2_| structure (Fig. 4 C). There are broad bands of second-order susceptibility in the lower left and upper right corners of the matrix centered on the anti-diagonal where *f*_1_ + *f*_2_ matches *f*_BaseCorrected_. After burst-spike removal, these bands of elevated second-order susceptibility disappear almost completely (Fig. 4 D) leaving only faint remainders of the broad bands when *f*_1_+*f*_2_ matches *f*_BaseCorrected_. Position and intensity of increased |*χ*_2_| in the burst-corrected case are very similar to the non-bursty low-CV P-unit shown in Fig. 3A. The projection of the anti-diagonals of the |*χ*_2_| matrix onto the main diagonal highlights the differences between original and burst-corrected response (Fig. 4 E). Bursts seem to increase the second-order susceptibility at the frequencies related to *f*_BaseCorrected_ but not the original *f*_Base_ (Fig. 4 C). Often, there are also peaks when *f*_1_+*f*_2_ matches *f*_EOD_ or *f*_EOD_ - *f*_BaseCorrected_ (Fig. 4 C-E).

### Adding bursts in the model revealed that the burst-induced increase of the second-order susceptibility is mediated not by the timing of the first spikes but by the number of spikes in a burst package

The number of spikes per burst varies within the same neuron (Fig. 4 A) and across cells (described by the so-called burst fraction, Fig. 7 A_iii,iv_). In the following, we address the influence of the spike number in a burst package on nonlinearity in a low-CV P-unit model (see methods) by systematically varying the size of the burst packages by adding one, two, or three successive spikes that occur with ISIs of exactly one EOD period. The original model response to a very weak amplitude modulation of the EOD carrier (red line in Fig. 5 A) is shown in the first row of the spike trains in Fig. 5 A. To mimic bursting, additional spikes were added with EOD-period ISIs. We applied the noise-splitting method (Novikov, 1965; Furutsu, 1963) (see methods and our previous work (Barayeu et al., 2024) for a detailed discussion) to uncover the full non-linear structure. As expected, the structure of elevated second-order susceptibility is related to the baseline firing rate *f*_Base_ of the model and its first harmonic (yellow lines of |*χ*_2_|, Fig. 5 B, top). The corresponding projection onto the main diagonal (Fig. 5 B, bottom) shows distinct peaks when *f*_1_ + *f*_2_ matches *f*_Base_, 2*f*_Base_ as well as a third peak at *f*_EOD_*/*2 *− f*_Base_. Adding exactly one burst spike already strongly changes the second-order susceptibility matrix. The bands of nonlinear susceptibility are much broader but even though the baseline firing rate is now doubled, the nonlinear structure still depends on *f*_BaseCorrected_ and its harmonic (pink edges, Fig. 5 C). On the other hand, the |*χ*_2_| values are reduced when *f*_1_ + *f*_2_ roughly matches *f*_EOD_*/*2 (orange dashed line Fig. 5 C). Increasing the number of spikes to three (Fig. 5 D) or four spikes (Fig. 5 E) adds more bands in the second-order susceptibility and introduces several elevations and dips in the projected diagonals. Although with more spikes per burst package the overall nonlinearity increases (more yellow in Fig. 5 E than in Fig. 5 B), some frequencies are enhanced but others are attenuated. The peak at *f*_BaseCorrected_ in the projected diagonal is increased with more spikes in a burst package (compare Fig. 5 B and Fig. 5 C–E). The frequencies on the vertical and horizontal lines at *f*_1_ = *f*_BaseCorrected_ and *f*_2_ = *f*_BaseCorrected_ in the second-order susceptibility are enhanced or attenuated depending on their position in the antidiagonal bands induced by bursts. Adding bursts in the model revealed that the burst-induced increase of the second-order susceptibility is mediated not by the timing of the first spikes but by the number of spikes in a burst package.

**Figure 5:**
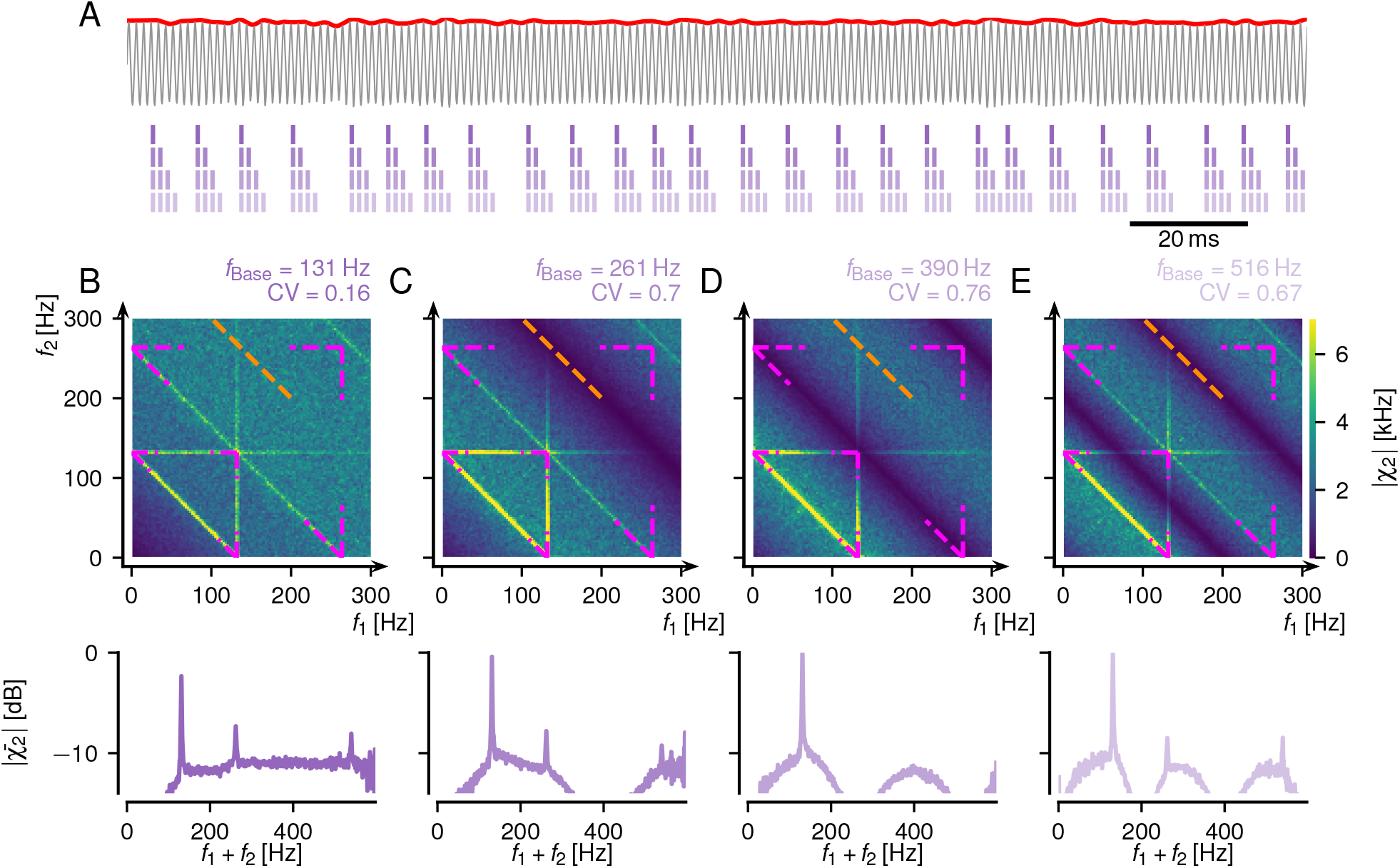
Adding spikes to a burst package increases the second-order susceptibility in low-CV models with 1 million stimulus realizations (see table 1 for model parameters of 2013-01-08-aa). A model with an intrinsic noise split (see methods). **A** Top: EOD carrier (gray) with a weak random amplitude modulation (RAM) stimulus (red). Bottom: spike trains with an increasing number of spikes per bust package. **D–E** Top: Absolute value of the second-order susceptibility. Bottom: Projected diagonal (see methods). **B** Original model 2013-01-08-aa. **C** Same as before but with two spikes per burst packages: The additional spike was added with an ISI of one EOD period following each spike in the original response. Pink lines – edges of the structure that occur when *f*_1_, *f*_2_, or *f*_1_ + *f*_2_ are equal to *f*_Base_ or its harmonic. Orange line when *f*_1_ + *f*_2_ is equal to half *f*_EOD_. **D** Same for three-spike burst packages. The only difference is that the pink structure refers to *f*_BaseCorrected_. **E** Same for four-spike burst packages.

### P-units of *Eigenmannia virescens* implement nonlinearity with very low-CVs

Bursting activity has been described for P-units of *Apteronotus leptorhynchus* (Chacron et al., 2004) and we have seen here, that busting increases and broadens their second-order susceptibility. The P-units of a related species, *Eigenmannia virescens*, however, do not show bursting activity in their firing pattern. Even in cases of high firing rates, there are no burst packages that are separated from each other by intervals of quiescence (Fig. 6 A_ii_ B_ii_). P-units of *Eigenmannia virescens* have generally lower CVs than the P-units of *Apteronotus leptorhynchus* (compare top marginal distributions in Fig. 7 A_i_ and B_i_). Nevertheless, the second-order susceptibility of both cells shown in Fig. 6 resembles that of bursty *Apteronotus leptorhynchus* P-units with broad bands of |*χ*_2_| values around *f*_Base_ (pink edges in Fig. 6 A_iii_ and B_iii_). There is also a fine line of non-linearity when *f*_1_ + *f*_2_ matches *f*_EOD_. *Eigenmannia virescens* have lower EOD frequencies than *Apteronotus leptorhynchus* and the *f*_Base_ of P-units can be as high as the *f*_EOD_ which is generally not the case in *Apteronotus leptorhynchus* in which it is usually below *f*_EOD_*/*2. In most P-units *f*_Base_ is close to *f*_EOD_*/*2, with very pronounced nonlinearity bands at *f*_1_ + *f*_2_ = *f*_Base_ and *f*_1_ + *f*_2_ = *f*_*EOD*_*/*2 (Fig. 6 A_iv_).

**Table 1:** Model parameters of LIF models.

| *cell* | $\beta$ | $\tau_m/\text{ms}$ | $\mu$ | $D/\text{ms}$ | $\tau_A/\text{ms}$ | $\Delta_A$ | $\tau_d/\text{ms}$ | $t_{ref}/\text{ms}$ |
| --- | --- | --- | --- | --- | --- | --- | --- | --- |
| 2013-01-08-aa | 4.5 | 1.20 | 0.59 | 0.001 | 37.52 | 0.01 | 1.18 | 0.38 |

**Figure 6:**
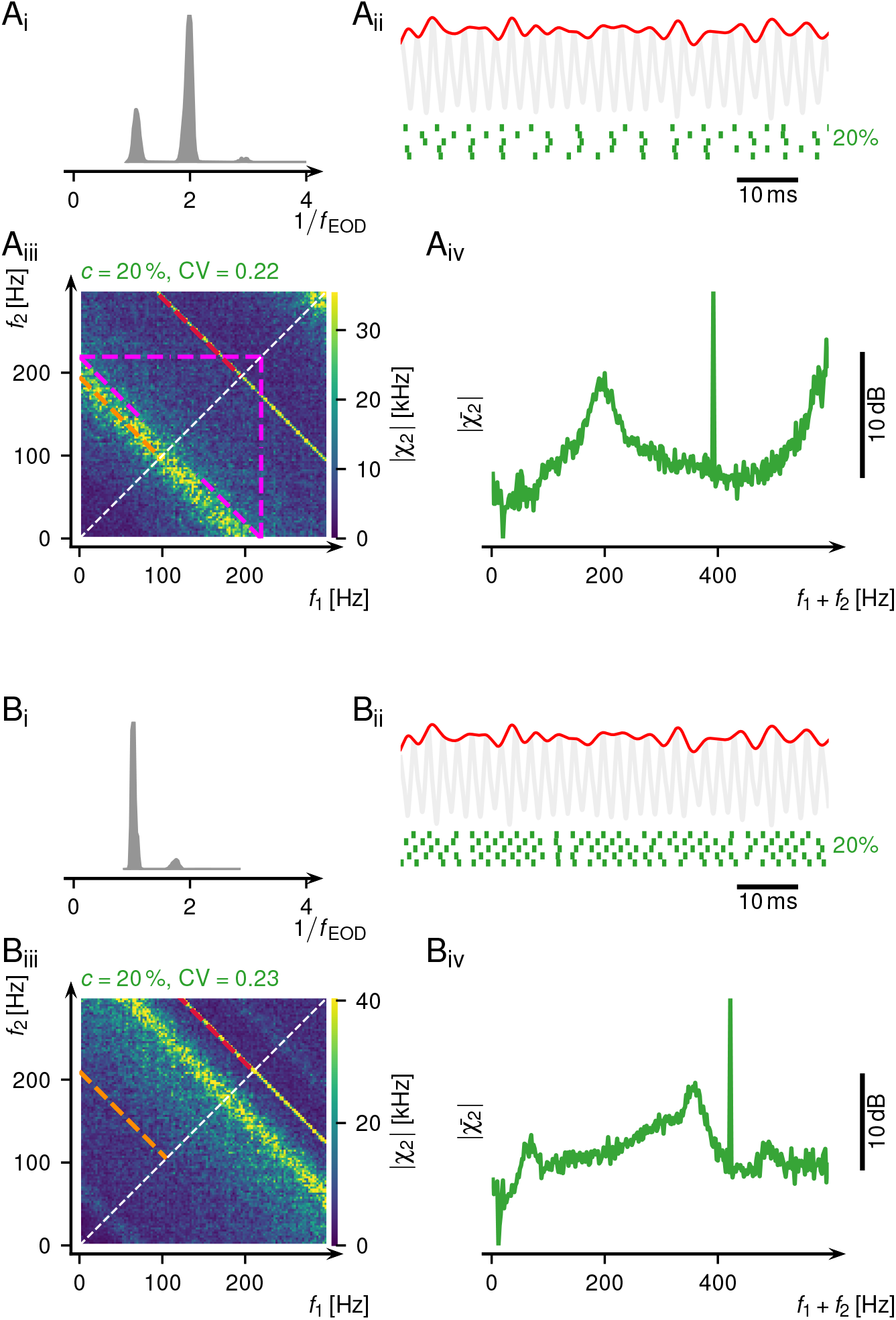
Response of experimentally measured P-units of *Eigenmannia virescens*. **A** Low-CV P-unit with strong second-order susceptibility when *f*_1_ + *f*_2_ = *f*_Base_. **A**_i_ Left: ISI distribution of the baseline activity. **A**_ii_ Top: EOD carrier (gray) with a RAM stimulus (red). Bottom: Spike raster of five consecutive stimulus presentations. **A**_iii_ Absolute value of the second-order susceptibility. Pink lines highlight when *f*_1_, *f*_2_, or *f*_1_ + *f*_2_ are equal to *f*_Base_. Orange line highlights the situation that *f*_1_ + *f*_2_ is equal to half *f*_EOD_. Red line highlights when *f*_1_ + *f*_2_ is equal to *f*_EOD_. **A**_iv_ Projected diagonal, calculated as the mean of the anti-diagonals of the matrix in A_iv_. **B** Same as A but for a low-CV P-unit with strong second-order susceptibility when *f*_1_ +*f*_2_ = *f*_Base_ (pink) and *f*_1_ +*f*_2_ = *f*_*EOD*_*/*2 (orange, in B_iii_).

**Figure 7:**
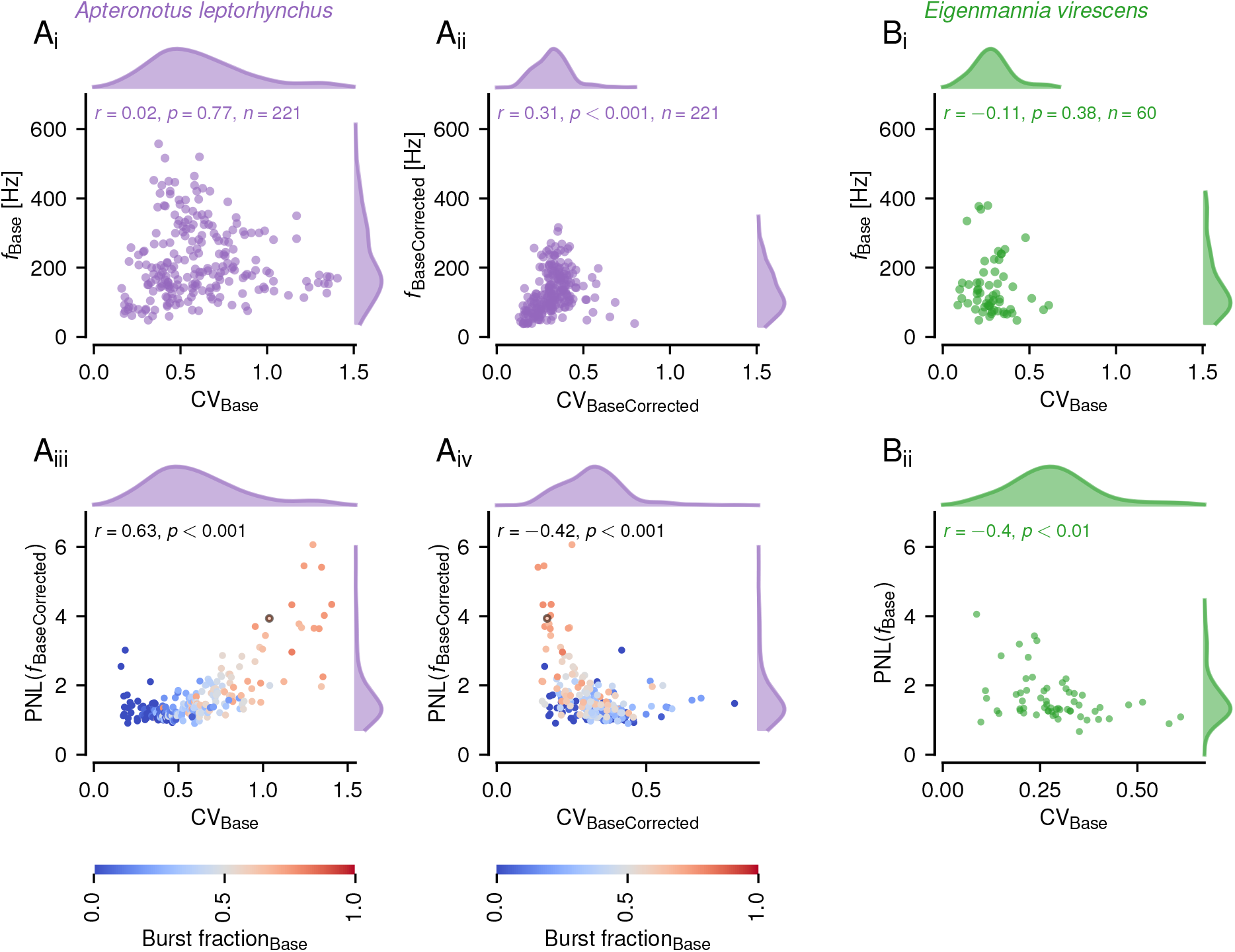
Population statistics of P-units in *Apteronotus leptorhynchus* (A, purple) and *Eigenmannia virescens* (B, green). *f*_Base_ – baseline firing rate before burst correction. *f*_BaseCorrected_ – baseline firing rate after burst correction. CV_Base_ – baseline CV before burst correction. CV_BaseCorrected_ – baseline CV after burst correction. **A**_i_, **A**_ii_, **B**_i_ Baseline statistics. Burst correction leads to a reduction of baseline CVs and mean firing rates in *Apteronotus leptorhynchus* (A_i,ii_). *Eigenmannia virescens* has lower CVs and firing rates during baseline (B_i_). **A**_iii_ Nonlinearity is strongly correlated with the CV. **A**_iv_ Gray circle corresponds to the cell in Fig. 4 B. The nonlinearity of cells with a high burst fractions is different if it is considered at *f*_Base_ and *f*_BaseCorrected_. **B**_ii_ The low-CV P-units of *Eigenmannia virescens* have pronounced nonlinearities.

### Bursts influence nonlinearity on a population level

So far we have illustrated the nonlinear encoding with single-cell examples, only. In the course of this study, we analyzed the responses of 282 P-units recorded over the past 15 years. 221 of these were recorded in *Apteronotus leptorhynchus* 60 cells in *Eigenmannia virescens*. The P-units in both species serve the same purpose and encode amplitude modulations of the self-generated EOD. In both species, P-units fire phase-locked to the EOD carrier, the probability of spiking, however depends on the amplitude of the carrier. The median of the average firing rates observed during the spontaneous, baseline, activity is similar while the distribution is wider in *A. leptorhynchus*. The distribution of the CVs of the ISI during spontaneous activity is much broader in *A. leptorhynchus* than in *E. virescens* (marginal distributions in Fig. 7 A_i_, B_i_). A statistically significant correlation between CV_Base_ and *f*_Base_ could not be found in either species. The high CVs, and high baseline rates, in *A. leptorhynchus* are attributable to the bursting response mode. When performing the burst correction, the distributions of *f*_BaseCorrected_ and CV_BaseCorrected_ are very similar in both species (compare Fig. 7 A_ii_ and B_i_). After burst correction, there is a statistically significant correlation between the corrected baseline firing rate and the corrected CV in *A. leptorhynchus*. The burst fraction, i.e. the fraction of ISIs that are part of bursts, correlates strongly with the baseline CV_Base_. Our previous work suggested an inverse correlation of the baseline CV_Base_ and the nonlinear encoding (Barayeu et al., 2024). We used the PNL(*f*_Base_) score to characterize the peakedness of the peak at *f*_BaseCorrected_ in the projected diagonal (Eq. (11)). We found previously that low-CV non-bursty P-units show higher levels of second-order susceptibility. Here, these cells are shown at the left border of Fig. 7 A_iii_. With higher burst fractions and thus a higher CV_Base_, we find that the nonlinear encoding increases statistically significantly (*r* = 0.63, *p <* 1%) in *A. leptorhynchus*. In *E. virescens*, however, there are similar levels of nonlinearity as in the burst-corrected CV condition (Fig. 7 A_iv_) despite the fact that there are no bursts (Fig. 7 B_ii_).

### P-units use bursting to encode stimulus amplitude

In the stimulus-driven case, most cells increase bursting when driven strongly (high response modulation, red markers are above and blue markers around the equality line, Fig. 8 A). After burst correction the mutual information is decreasing for cells with a high burst fraction (Fig. 8 B). These findings highlight that some P-units use bursting as a mechanism to encode a wide dynamic range of contrasts. Cells that already possess high burst fractions during baseline cannot increase their firing even further (high burst fractions values close to the equality line, Fig. 8 A) and thus cannot use bursting as a mechanism for contrast encoding.

**Figure 8:**
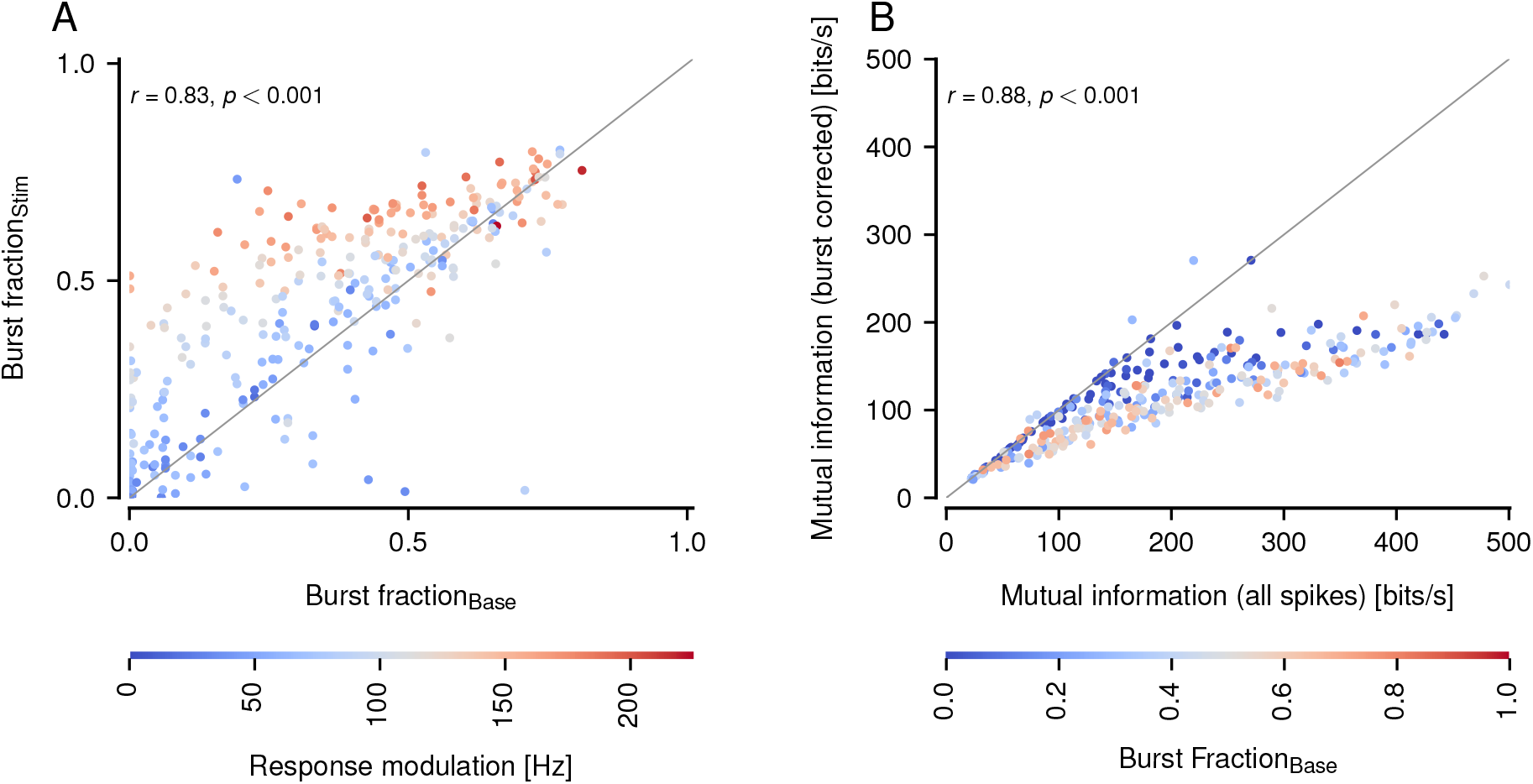
Bursting contributes to stimulus encoding in all P-units. **A** High bust fraction cells do not increase their bursting during stimulation and are close to the gray equality line. Cells that are not intrinsic bursters, with a low burst fraction, increase their bursting with stronger stimulus amplitudes (red colors). **B** The mutual information after burst correction is lower compared to the mutual information of all spikes before burst correction.

## Discussion

The classic view of bursting is, that bursts of action potentials are employed to mark the occurrence of special features in the stimulus and then, with a high signal-to-noise ratio, relay this information faithfully to higher processing stages (Krahe and Gabbiani, 2004; Oswald et al., 2004; Gabbiani et al., 1996). In the primary electroreceptor afferents recorded here, it is long known that a fraction of P-units shows bursting even in their spontaneous activity (Bastian, 1981; Chacron et al., 2004; Krahe and Gabbiani, 2004; Metzen et al., 2016). Slice recordings showed that the ISI statistics of P-units are transported to higher-order neurons, the pyramidal cells in the electrosensory line lobe (ELL, Khanbabaie et al., 2010). Bursting in the primary sensory afferents might thus have an impact on higher-order cells. During spontaneous activity, however, when there are no features to highlight, bursting appears like a waste of energy and a source of noise for downstream processing. This has led to the surmise that these cells might have been damaged by the recording electrode and are in fact unhealthy. Modeling work, however, concluded that tonically firing and bursting neurons might serve different purposes. The first are better suited to encode the time course of the stimulus while the latter are better suited for feature detection (Chacron et al., 2001b, 2004). With this account, and also the analyses presented in the companion paper (Schlungbaum et al., 2024), we provide further experimental evidence hinting towards a functional role of bursting in P-units.

The cellular mechanism leading to bursting in P-units is so far unknown. Chacron and colleagues modeled it as a spike-triggered exciting current in conjunction with an adaptive spiking threshold that induces bursts and also terminates burst packages (Chacron et al., 2001b, 2004). Their conclusion that tonically firing and bursting P-units serve strictly different purposes is in some ways in conflict with the data shown in Fig. 4 B and Fig. 8 B. After burst-correction, i.e. the removal of the burst spikes while keeping the first spike of each burst package, the linear susceptibility of the neuron was strongly reduced. If the loss was just a consequence of reduced firing rates (Schneidman et al., 2000; Borst and Haag, 2001) we would expect the reduction to be similar for all frequency components. It seems, that this is not the case. Rather, the low-frequency components are reduced more strongly than mid-or high-frequency components. Further, if the occurrence of bursts would not carry stimulus-related information the mutual information rate estimated from the stimulus-response coherence, a linear measure, would not suffer from burst spike removal as is observed in Fig. 8 B, burst spikes that are mere additions unrelated to the stimulus would act as noise and reduce the information content of the responses. Rather, those cells that show bursting also under baseline conditions are those that lose most information content when the burst spikes are removed under stimulation. Detailed modeling and theoretical analyses are presented in the companion paper (Schlungbaum et al., 2024) and also suggest that bursts contribute to linear encoding. Usually, bursts in P-units have been discussed considering their contribution to nonlinearity (Chacron et al., 2004) with their impact on linear encoding being mainly neglected. The increase of linearity via bursts is not universal since in other cells, as in pyramidal cells of the ELL, the linear encoding of the stimulus is attributed to isolated spikes and not to bursts (Oswald et al., 2004).

The focus of this work is, however, on the effect of bursting on nonlinear encoding. In our previous work, we described that low-CV non-bursting P-units show non-linear susceptibility under certain stimulus conditions, an effect not seen in high-CV non-bursty P-units (Barayeu et al., 2024). These nonlinear interactions were known for pure LIF models (Voronenko and Lindner, 2017) but were not described for cells that respond to amplitude modulations of a carrier signal, i.e. the fish’s own electric field. The CV is interpreted as a proxy of the intrinsic noise which is a simplification of the real situation (Vilela and Lindner, 2009). In this line of thinking, it is surprising that bursting neurons that show a very high CV in their baseline activity actually have strong nonlinear susceptibilities, exceeding the ones observed in non-bursty low-CV P-units (Fig. 7 A_iv_). While the non-bursty low-CV cells show nonlinear susceptibility under very specific stimulus conditions, the bursty P-units show nonlinear encoding in much wider ranges. Two nonlinearity structures were present in the second-order susceptibility: the wide burst-induced bands and the sharp bands at *f*_BaseCorrected_ (Fig. 4 C), that were enhanced by bursts (Fig. 5 C–E). Using the P-unit LIF model we found that increased nonlinearity is not driven by the timing of the first spike in the burst package but is defined by the number of spikes in a burst package, with more burst spikes inducing higher nonlinearity. In pyramidal cells of the ELL, it was demonstrated that bursting improves feature detection by improving the signal-to-noise ratio (Oswald et al., 2004; Gabbiani et al., 1996). An improved signal-to-noise ratio and sharpened tuning curves have been associated with bursts in the auditory system (Eggermont and Smith, 1996) possibly due to a more reliable synaptic transmission (Lisman, 1997; Csicsvari et al., 1998) induced by bursts. As bursts contribute to linear and nonlinear stimulus encoding, reading out all information contained in the responses will indeed require a nonlinear decoder (Chacron et al., 2004).

We compared the second-order susceptibility of P-units in *Apteronotus leptorhynchus* with their counterparts recorded in *Eigenmannia virescens*, a closely related species. The basic response properties are similar, but we do not observe bursting. Surprisingly, nonlinear stimulus susceptibility is similar to those of bursting P-units in *A. leptorhynchus*, despite the lack of bursting. Does this imply that these cells are different? A similar question arises when we compare bursting and nonbursting P-units within *A. leptorhynchus*, are these indeed different versions of P-units? Removing burst spikes reduces the CV of bursting P-units and makes them appear as non-bursty cells. Accordingly, the nonlinear susceptibility is reduced to very similar levels (compare Fig. 4). It seems conceivable, that these cells are actually the same but bursting P-units have a very high sensitivity and their own EOD is enough to provoke high-frequency firing until the adaptation currents terminate the barrage of action potentials. In *E. virescens* the EOD-frequencies are in a different range (300 – 450 Hz as compared to 600 – 1000 Hz in *A. leptorhynchus* (Henninger et al., 2020)), that is, the minimal interspike interval is at least twice as long as in *A. leptorhynchus* which may have an effect on the dynamics of the adaptation currents that accumulate enough for P-units in *A. leptorhynchus* but not in *E. virescens*.

### Functional role of bursting and nonlinear susceptibility

Bursting has previously been associated with an increased second-order susceptibility in ampullary cells of paddlefish facilitating prey detection (Neiman and Russell, 2011), to encode different stimulus strengths as e.g. temperature changes (Longtin and Hinzer, 1996), or muscle stretch intensity (Birmingham et al., 1999). In the context of the wave-type weakly electric fish the interference patterns experienced during social encounters are important stimuli (Benda et al., 2005; Hupé and Lewis, 2008; Stamper et al., 2010; Savard et al., 2011; Henninger et al., 2018, 2020). Extracting such beats and higher-order envelopes (Stamper et al., 2012) requires nonlinear processes found in downstream neurons (Middleton et al., 2006, 2007). The encoding of social envelopes can also be found in P-units with stronger nonlinearities, lower firing rates, and higher CVs (Savard et al., 2011). In this account, high CVs were associated with increased bursting (Fig. 7 A_iii_) and higher nonlinear susceptibility. Thus, it is likely that the high-CV nonlinear envelope encoding P-units described by Savard et al. (2011) are bursty. In that case, the findings by (Savard et al., 2011) would be well in line with the data provided in this work.

In the real electrosensory world, white noise stimuli are rarely encountered, see however the prey signals for example experienced and encoded by ampullary electroreceptors of the paddlefish (Neiman and Russell, 2011). For P-units in *A. leptorhynchus* and *E. virescens* the beats arising from the interference of EODs with different frequencies are behaviorally relevant stimuli (Benda et al., 2005; Hupé and Lewis, 2008; Stamper et al., 2010; Savard et al., 2011; Henninger et al., 2018, 2020). In our previous work, we could show that the P-unit responses to stimulation with combinations of beats (e.g. when three fish meet) can be predicted from the second-order susceptibility estimated from white noise stimulation (Barayeu et al., 2024). Under the condition that the sum of the beat frequencies matches the baseline firing rate of the P-unit, a nonlinear interference occurs that induces responses at the sum of the individual frequencies, which can only arise through nonlinear processes. The occurrence of this nonlinear interaction peak in the response power spectra might even explain why the presence of a strong foreground signal can facilitate the detection of a weak signal (Schlungbaum and Lindner, 2023), a situation observed in recordings of animal interactions during courtship in freely moving animals (Henninger et al., 2018). In the previous work, we focused on low-CV P-units that showed this kind of nonlinear susceptibility at very specific stimulus situations (Barayeu et al., 2024). Bursts broaden the range of nonlinear encoding around the antidiagonal defined by the baseline rate of the neuron (Fig. 4). Interestingly, it is the burst-corrected but not the baseline firing rate including burst spikes, that predicts frequency combinations prone to nonliterary. In conjunction with a high degree of heterogeneity with respect to the baseline firing rates (Hladnik and Grewe, 2023), the electrosensory periphery seems to be well able to nonlinearly respond to a broad range of stimulus combinations.

### Nonlinearity necessary to sustain in different species?

In this chapter, it was demonstrated that similar nonlinearity values can be implemented with different mechanisms, namely with low-CV P-units in *Eigenmannia virescens* or bursting in *Apteronotus leptorhynchus* (Fig. 7 A_iv_, B_ii_). This suggests that nonlinearity might be an important feature, necessary to sustain in different species of weakly electric fish. If traits develop independently, with the trait not being present in a common ancestor, the process is termed convergent evolution. Examples of convergent evolution are blue eyes and light skin in humans (Edwards et al., 2010), the camera eye present in mammals, jellyfish, and squids (Kozmik et al., 2008) or the evolution of wave-type and pulse-type weakly electric fish (Lavoué et al., 2012). Slightly different traits in closely related species imply divergent evolution. An example of divergent evolution are dogs and wolfs that descend from the same ancestor (Vilà et al., 1999) or Darwin’s finches that developed beaks with different shapes (Ford et al., 1973). *Apteronotus leptorhynchus* and *Eigenmannia virescens* are closely related species belonging to the same superfamily Apteronotoidea, but to different families of Apteronotidae and Sternopygidae (Albert, 2001). The nonlinearity in the primary sensory afferents of more weakly electric fish species should be accessed in further studies, to increase the certainty if such nonlinearity might have been present in the common ancestor of *Eigenmannia virescens* and *Apteronotus leptorhynchus* and test whereas this sustained nonlinearity might be an example of divergent evolution.

Nonlinear encoding might thus be a feature of the cellular responses that in some way is critical for sensory encoding and thus needs to be maintained by one or the other mechanism. The way how it is realized, however, differs between species.

## Methods

The following descriptions of the experimental procedures and analyses are the same as in our companion paper (Barayeu et al., 2024) deviations from the original descriptions are highlighted.

### Experimental subjects and procedures

Within this project we re-evaluated datasets that were recorded between 2010 and 2023 at the Ludwig Maximilian University (LMU) München and the Eberhard-Karls University Tübingen. All experimental protocols complied with national and European law and were approved by the respective Ethics Committees of the Ludwig-Maximilians Universität München (permit no. 55.2-1-54-2531-135-09) and the Eberhard-Karls Unversität Tübingen (permit no. ZP 1/13 and ZP 1/16). The final sample consisted of 221 P-units recorded in 71 weakly electric fish of the species *Apteronotus leptorhynchus* and, in addition to the analyses in the companion paper, 60 P-units recorded in 17 *Eigenmannia virescens*. The original electrophysiological recordings were performed on male and female fish obtained from a commercial supplier for tropical fish (Aquarium Glaser GmbH, Rodgau, Germany). The fish were kept in tanks with a water temperature of 25 ± 1 °C and a conductivity of around 270 µS*/*cm under a 12 h:12 h light-dark cycle.

Before surgery, the animals were deeply anesthetized via bath application with a solution of MS222 (120 mgample to fire an action potential, the membrane voltage must exceed a threshold nonlinearity. Even though, linear encoding schemes are commonly used and often successfully describe large parts of sensory encoding nonlinear mechanisms such as thresholds and saturations are well known to be crucial to encode behaviorally relevant features in the stimulus space not captured by linear methods. Here we analyplication of liquid lidocaine hydrochloride (20 mg/ml, bela-pharm GmbH). During the surgery water supply was ensured by a mouthpiece to maintain anesthesia with a solution of MS222 (100 mg/l) buffered with Sodium Bicarbonate (100 mg/l). After surgery, fish were immobilized by intramuscular injection of from 25 µl to 50 µl of tubocurarine (5 mg/ml dissolved in fish saline; Sigma-Aldrich). Respiration was then switched to normal tank water and the fish was transferred to the experimental tank.

### Experimental setup

For the recordings fish were positioned centrally in the experimental tank, with the major parts of their body submerged into the water. Those body parts that were above the water surface were covered with paper tissue to avoid drying of the skin. Local analgesia was refreshed in intervals of two hours by cutaneous reapplication of Lidocaine (2 %; bela-pharm, Vechta, Germany) around the surgical wounds. Electrodes (borosilicate; 1.5 mm outer diameter; GB150F-8P; Science Products, Hofheim, Germany) were pulled to a resistance of 50–100 MΩ (model P-97; Sutter Instrument, Novato, CA) and filled with 1 M KCl solution. Electrodes were fixed in a microdrive (Luigs-Neumann, Ratingen, Germany) and lowered into the nerve (Fig. 9, blue triangle). Recordings of electroreceptor afferents were amplified and lowpass filtered at 10 kHz (SEC-05, npi-electronics, Tamm, Germany, operated in bridge mode). All signals, neuronal recordings, recorded EOD and the generated stimulus, were digitized with sampling rates of 20 or 40 kHz (PCI-6229, National Instruments, Austin, TX). RELACS (www.relacs.net) running on a Linux computer was used for online spike and EOD detection, stimulus generation, and calibration. Recorded data was then stored on the hard drive for offline analysis.

**Figure 9:**
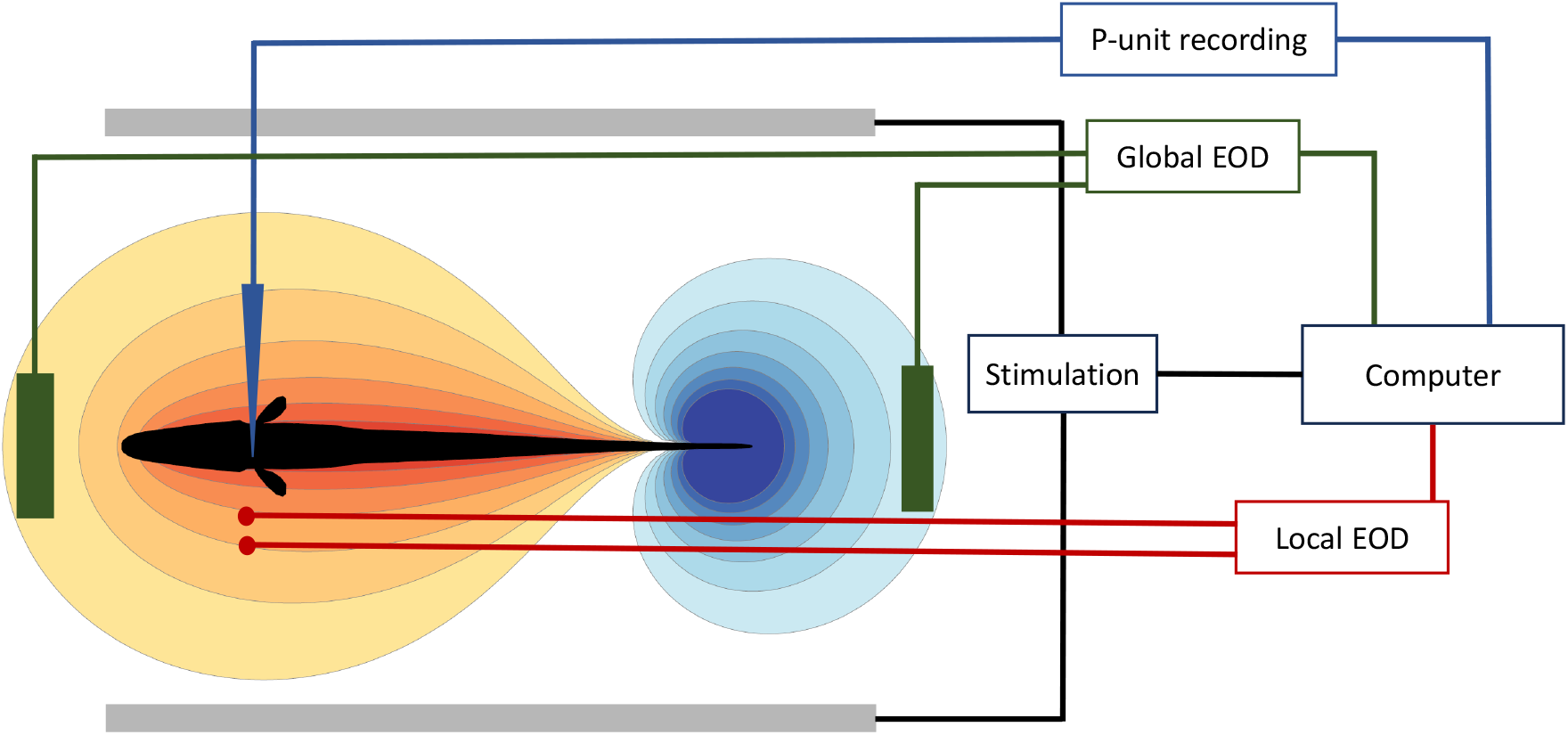
Electrophysiolocical recording setup. The fish, depicted as a black scheme and surrounded by isopotential lines, was positioned in the center of the tank. Blue triangle – electrophysiological recordings were conducted in the posterior anterior lateral line nerve (pALLN). Gray horizontal bars – electrodes for the stimulation. Green vertical bars – electrodes to measure the *global EOD* placed isopotential to the stimulus, i.e. recording fish’s unperturbed EOD. Red dots – electrodes to measure the *local EOD* picking up the combination of fish’s EOD and the stimulus. The local EOD was measured with a distance of 1 cm between the electrodes. All measured signals were amplified, filtered, and stored for offline analysis.

### Identification of P-units

The neurons were classified into cell types during the recording by the experimenter. P-units were classified based on baseline firing rates of 50–450 Hz and a clear phase-locking to the EOD and their responses to amplitude modulations of their own EOD (Grewe et al., 2017; Hladnik and Grewe, 2023).

### Electric field recordings

The electric field of the fish was recorded in two ways: 1. we measured the so-called *global EOD* with two vertical carbon rods (11 cm long, 8 mm diameter) in a head-tail configuration (Fig. 9, green bars). The electrodes were placed isopotential to the stimulus. This signal was differentially amplified with a factor between 100 and 500 (depending on the recorded animal) and band-pass filtered (3 to 1500 Hz pass-band, DPA2-FX; npi electronics, Tamm, Germany). 2. The so-called *local EOD* was measured with 1 cm-spaced silver wires located next to the left gill of the fish and orthogonal to the fish’s longitudinal body axis (amplification 100 to 500 times, band-pass filtered with 3 to 1 500 Hz pass-band, DPA2-FX; npi-electronics, Tamm, Germany, Fig. 9, red markers). This local measurement recorded the combination of the fish’s own field and the applied stimulus and thus serves as a proxy of the transdermal potential that drives the electroreceptors.

### Stimulation

The stimulus was isolated from the ground (ISO-02V, npi-electronics, Tamm, Germany) and delivered via two horizontal carbon rods (30 cm length, 8 mm diameter) located 15 cm laterally to the fish (Fig. 9, gray bars). The stimulus was calibrated with respect to the local EOD.

### White noise stimulation

The fish were stimulated with band-limited white noise stimuli with a cut-off frequency of 300 or 400 Hz. The stimulus intensity is given as the contrast, i.e. the standard deviation of the white noise stimulus in relation to the fish’s EOD amplitude. The contrast varied between 1 and 20 %. Only cell recordings with at least 10 s of white noise stimulation were included for the analysis. To create random amplitude modulations (RAM) for P-unit recordings, the EOD of the fish was multiplied with the desired random amplitude modulation profile (MXS-01M; npi electronics).

### Data analysis

Data analysis was performed with Python 3 using the packages matplotlib (Hunter, 2007), numpy (Walt et al., 2011), scipy (Virtanen et al., 2020), pandas (McKinney et al., 2010), nixio (Stoewer et al., 2014), and thunderfish (https://github.com/bendalab/thunderfish).

### Baseline analysis

The baseline firing rate *f*_Base_ was calculated as the number of spikes divided by the duration of the baseline recording (on average 18 s). The coefficient of variation (CV) was calculated as the standard deviation of the interspike intervals (ISI) divided by the average ISI: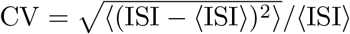. If the baseline was recorded several times in a recording, the average *f*_Base_ and CV were calculated.

### Characterization of bursting

In this account, we focus on the effect of bursting. Bursty cells are characterized by a bimodal ISI distribution with a strong peak close to the EOD period. The burstiness of the neuronal baseline response was described by the so-called burst fraction (Chacron et al., 2004; Metzen et al., 2016) which is the fraction of ISIs that are shorter than a certain threshold (usually 1.5 EOD periods). In some P-units, however, the burst threshold was adapted to best separate the bimodal distribution by visual inspection. In the final P-units sample, 149 cells had a burst threshold of 1.5 EOD periods, 59 cells had a burst threshold of 2.5 EOD periods, 7 cells had a burst threshold of 3.5 EOD periods, 4 cells had a burst threshold of 4.5 EOD periods and 1 cell had a burst threshold of 5.5 EOD periods. The burst fraction was calculated as the number of ISIs below the burst threshold divided by the total number of interspike intervals during baseline activity.

For cells that showed bursting, we additionally calculated the baseline firing rate after burst correction *f*_BaseCorrected_ and the baseline CV after burst correction CV_BaseCorrected_, by removing all spikes except for the first of a burst package.

### White noise analysis

In the stimulus driven case, the neuronal activity of the recorded cell is modulated around the average firing rate that is similar to *f*_Base_ and in that way encodes the time-course of the stimulus. The time-dependent response of the neuron was estimated from the spiking activity

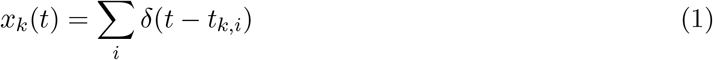

recorded for each stimulus presentation, *k*, by kernel convolution with a Gaussian kernel

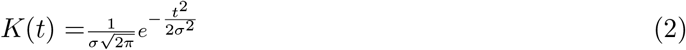

with *σ* the standard deviation of the Gaussian which was set to 2.5 ms if not stated otherwise. For each trial *k* the *x*_*k*_(*t*) is convolved with the kernel *K*(*t*)

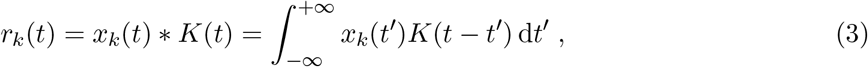

where *∗* denotes the convolution. *r*(*t*) is then calculated as the across-trial average

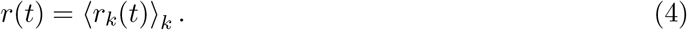

To quantify how strongly the neuron is driven by the stimulus we quantified the response modulation as the standard deviation 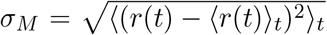, where ( · )_*t*_ indicates averaging over time.

### Spectral analysis

The neuron is driven by the stimulus and thus the spiking response *x*(*t*), Eq. (1), depends on the stimulus *s*(*t*). To investigate the relation between stimulus and response we calculated the first- and second-order susceptibility of the neuron to the stimulus in the frequency domain. The Fourier transforms of *s*(*t*) and *x*(*t*) are denoted as 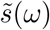 and 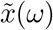 and were calculated according to 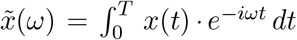 *dt*, with *T* being the signal duration. Stimuli had a duration of 10 s and spectra of stimulus and response were calculated in separate segments of 0.5 s with no overlap resulting in a spectral resolution of 2 Hz.

The power spectrum of the stimulus *s*(*t*) was calculated as

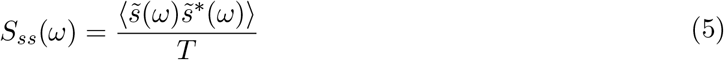

with 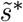 being the complex conjugate and ⟨…⟩ denoting averaging over the segments. The power spectrum of the spike trains *S*_*xx*_(*ω*) was calculated accordingly. The cross-spectrum *S*_*xs*_(*ω*) between stimulus and evoked spike trains was calculated according to

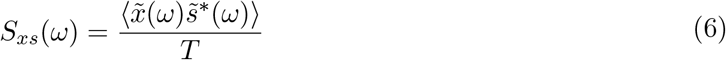

From *S*_*xs*_(*ω*) and *S*_*ss*_(*ω*) we calculated the linear susceptibility (transfer function) as

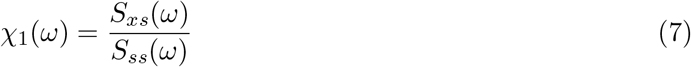

The second-order cross-spectrum that depends on the two frequencies *ω*_1_ and *ω*_2_ was calculated according to

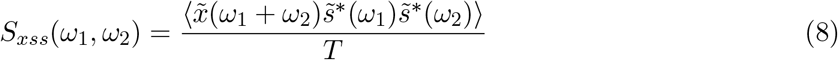

The second-order susceptibility was calculated by dividing the higher-order cross-spectrum by the spectral power at the respective frequencies.

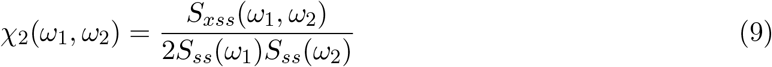

The absolute value of a second-order susceptibility matrix is visualized in Fig. 5. There the upper right quadrant characterizes the nonlinearity in the response *x*(*t*) at the sum frequency of the two input frequencies. The not shown lower right and upper left quadrants (as in Barayeu et al. (2024)) characterize the nonlinearity in the response *x*(*t*) at the difference of the input frequencies.

The coherence was calculated as

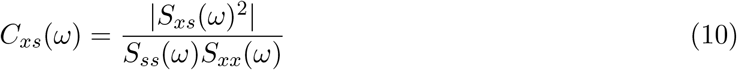

Mutual information was calculated as :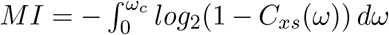, where *ω*_*c*_ is the cutoff frequency.

### Nonlinearity index

We expect to see nonlinear susceptibility when *ω*_1_ + *ω*_2_ = *f*_Base_. To characterize this we calculated the peakedness of the nonlinearity (PNL) as

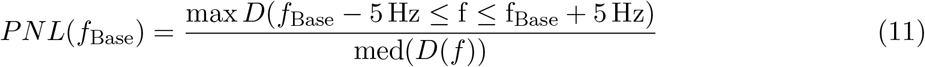

For this index, the second-order susceptibility matrix was projected onto the diagonal *D*(*f*), by taking the mean of the anti-diagonals. The peakedness at the frequency *f*_Base_ in *D*(*f*) was quantified by finding the maximum of *D*(*f*) in the range *f*_Base_ *±* 5 Hz (Fig. 4 E) and dividing it by the median of *D*(*f*).

If the same frozen noise was recorded several times in a cell, each noise repetition resulted in a separate second-order susceptibility matrix. The mean of the corresponding PNL(*f*_Base_) values was used for the population statistics in Fig. 7.

### Leaky integrate-and-fire models

Leaky integrate-and-fire (LIF) models with a carrier were constructed to reproduce the specific firing properties of P-units (Chacron et al., 2001a; Sinz et al., 2020). The sole driving input into the P-unit model during baseline, i.e. when no external stimulus was given, is the fish’s own EOD modeled as a cosine wave

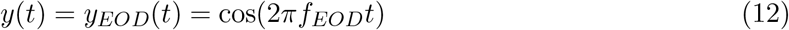

with the EOD frequency *f*_*EOD*_ and an amplitude normalized to one.

In the model, the input *y*(*t*) was then first thresholded to model the synapse between the primary receptor cells and the afferent.

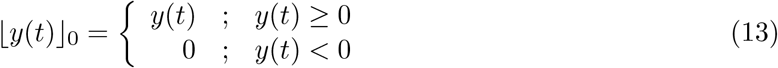

*ly*(*t*)*J*_0_ denotes the threshold operation that sets negative values to zero (Fig. 10 A).

**Figure 10:**
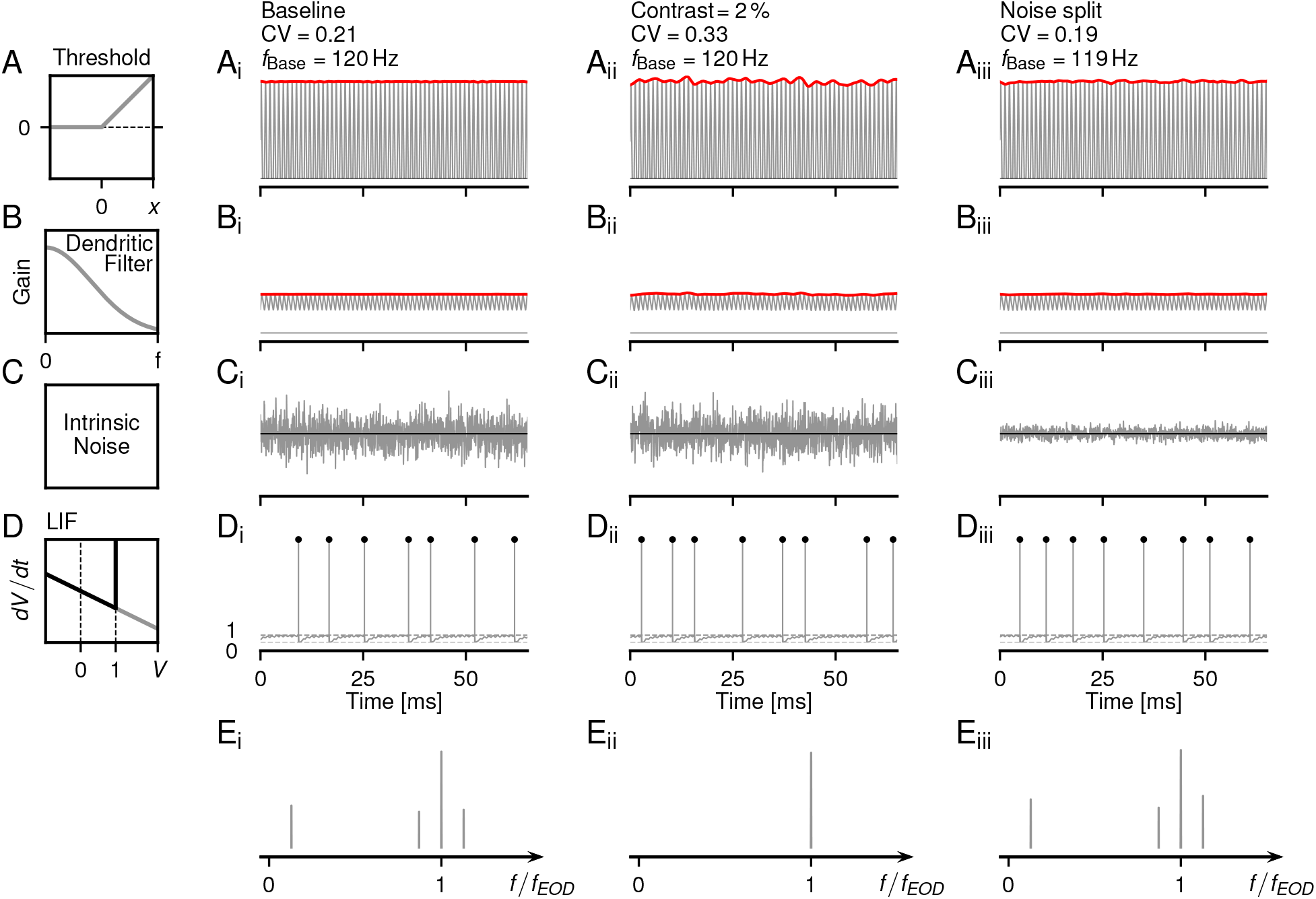
Flowchart of a LIF P-unit model with EOD carrier. The main steps of the model are depicted in the left model column (Model cell identifier 2012-07-03-ak, see table 1 for model parameters). The three other columns show the relevant signals in three different settings. (i) the baseline situation, no external stimulus, only the animal’s self-generated EOD (i.e. the carrier) is present (ii) RAM stimulation, the carrier is amplitude modulated with a weak (2 % contrast) stimulus, (iii) Noise split condition in which 90 % of the internal noise is used as a driving stimulus scaled with the correction factor *ρ* (see text). Note: that the firing rate and the CV of the ISI distribution is the same in this and the baseline condition. **A** Thresholding: a simple linear threshold was applied to the EOD carrier, Eq. (12). The red line on top depicts the amplitude modulation (AM). **B** Dendritic low-pass filtering attenuates the carrier. **C** A Gaussian noise is added to the signal in B. Note the reduced internal noise amplitude in the noise split (iii) condition. **D** Spiking output of the LIF model in response to the addition of B and C. **E** Power spectra of the LIF neuron’s spiking activity. Under the baseline condition (E_i_) there are several peaks, from left to right, at the baseline firing rate *f*_Base_, *f*_*EOD*_ *− f*_Base_ *f*_*EOD*_, and *f*_*EOD*_ + *f*_Base_. In the stimulus-driven regime (E_ii_), there is only a peak at *f*_EOD_, while under the noise split condition (E_iii_) again all peaks are present.

**Figure 11:**
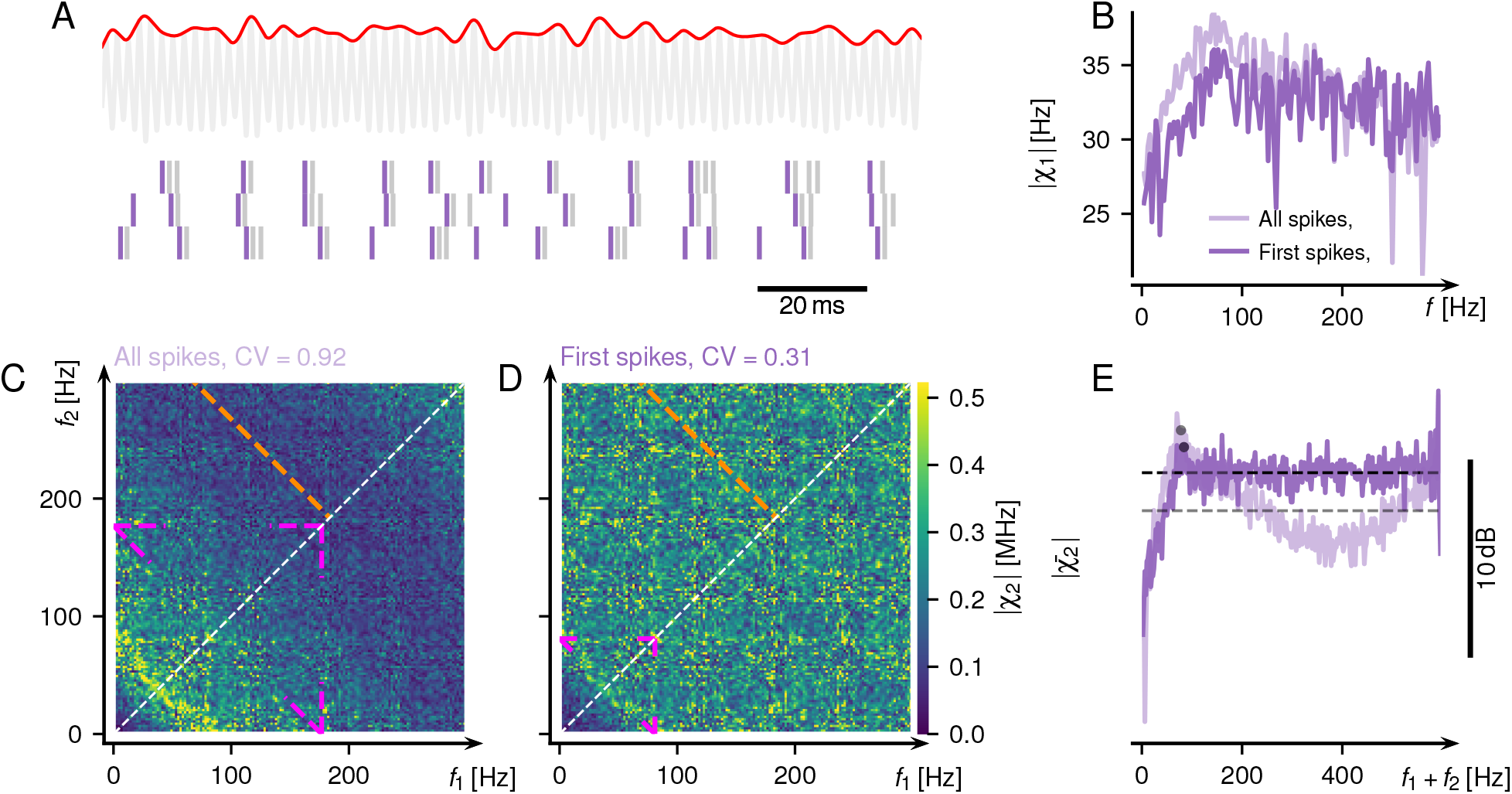
Response of two experimentally measured bursty P-units before and after burst correction (A, B). Burst correction has three effects: 1) It lowers the baseline frequency from *f*_Base_ to *f*_BaseCorrected_. 2) It decreases the overall second-order susceptibility. 3) It decreases the size of the *f*_BaseCorrected_ peak in the projected diagonal. **A**_i_ Left: ISI distribution during baseline. Right: Baseline power spectrum of the firing rate. **A**_ii_ Top: EOD carrier (gray) with RAM (red). Bottom: Dark purple – spikes after burst correction. **A**_iii_ First-order susceptibility (Eq. (7)). Light purple – based on all spikes (before burst correction). Dark purple – based only on the first spike of a burst package (after burst correction). **A**_iv_ Absolute value of the second-order susceptibility without burst correction. Pink lines – edges of the structure when *f*_1_, *f*_2_ or *f*_1_ + *f*_2_ are equal to *f*_Base_. Orange line part of the structure when *f*_1_ + *f*_2_ is equal to half *f*_EOD_. **A**_v_ Absolut value of the second-order susceptibility after burst correction. Pink lines – edges of the structure when *f*_1_, *f*_2_ or *f*_1_ + *f*_2_ are equal to *f*_BaseCorrected_. Orange line as in A_iv_. **A**_vi_ Projected diagonals, calculated as the mean of the anti-diagonals in the matrices in A_iv,v_. Colors as in A_iii_. Gray marker – *f*_BaseCorrected_. Dashed lines – median of the projected diagonals.

The resulting receptor signal was then low-pass filtered to approximate passive signal conduction in the afferent’s dendrite (Fig. 10 B)

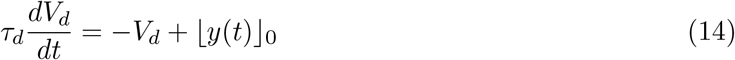

with *τ*_*d*_ as the dendritic time constant. Dendritic low-pass filtering was necessary to reproduce the loose coupling of P-unit spikes to the EOD while maintaining high sensitivity at small amplitude modulations. Because the input was dimensionless, the dendritic voltage was dimensionless, too. The combination of threshold and low-pass filtering extracts the amplitude modulation of the input *y*(*t*).

The dendritic voltage *V*_*d*_(*t*) was the input to a leaky integrate-and-fire (LIF) model

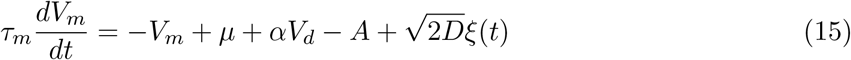

where *τ*_*m*_ is the membrane time-constant, *µ* is a fixed bias current, *α* is a scaling factor for *V*_*d*_, *A* is an inhibiting adaptation current, and 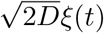 is a white noise with strength *D*. All state variables except *τ*_*m*_ are dimensionless.

The adaptation current *A* followed

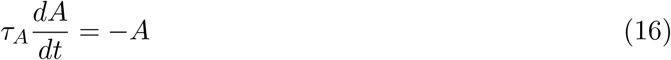

with adaptation time constant *τ*_*A*_.

Whenever the membrane voltage *V*_*m*_(*t*) crossed the spiking threshold *θ* = 1 a spike was generated, *V*_*m*_(*t*) was reset to 0, the adaptation current was incremented by Δ*A*, and integration of *V*_*m*_(*t*) was paused for the duration of a refractory period *t*_*ref*_ (Fig. 10 D).

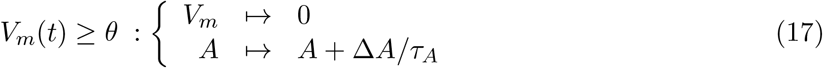

### Numerical implementation

The model’s ODEs were integrated by the Euler forward method with a time-step of Δ*t* = 0.05 ms. The intrinsic noise *ξ*(*t*) (Eq. (15), Fig. 10 C) was added by drawing a random number from a normal distribution *𝒩* (0, 1) with zero mean and standard deviation of one in each time step *i*. This number was multiplied with 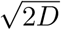 and divided by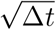 :

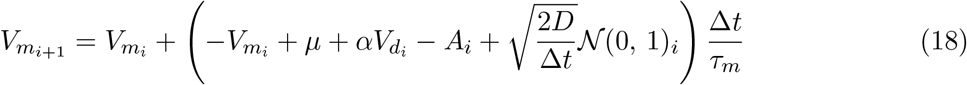

### Model parameters

The eight free parameters of the P-unit model *β, τ*_*m*_, *µ, D, τ*_*A*_, Δ_*A*_, *τ*_*d*_, and *t*_*ref*_, were fitted to both the baseline activity (baseline firing rate, CV of ISIs, serial correlation of ISIs at lag one, and vector strength of spike coupling to EOD) and the responses to step-like increases and decreases in EOD amplitude (onset-state and steady-state responses, effective adaptation time constant). For each simulation, the start parameters *A, V*_*d*_ and *V*_*m*_ were drawn from a random starting value distribution, estimated from a 100 s baseline simulation after an initial 100 s of simulation that was discarded as a transient.

### Stimuli for the model

The model neurons were driven with similar stimuli as the real neurons in the original experiments. To mimic the interaction with one or two foreign animals the receiving fish’s EOD (Eq. (12)) was normalized to an amplitude of one and the respective EODs of a second or third fish were added.

The random amplitude modulation (RAM) input to the model was created by drawing random amplitude and phases from Gaussian distributions for each frequency component in the range 0–300 Hz. An inverse Fourier transform was applied to get the final amplitude RAM time course. The input to the model was then

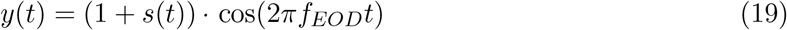

From each simulation run, the first second was discarded and the analysis was based on the last second of the data. The resulting spectra thus have a spectral resolution of 1 Hz.

### Model noise split into a noise and a stimulus component

According to previous works (Lindner, 2022) the total noise of a LIF model (*ξ*) can be split up into several independent noise processes. Here we split the internal noise into two parts: (i) One part is treated as a driving input signal *s*_*ξ*_(*t*), a RAM stimulus where frequencies above 300 Hz are discarded (Eq. (20)), and used to calculate the cross-spectra in Eq. (8) and (ii) the remaining noise 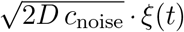 that is treated as pure noise (Eq. (22)). In this way the effective signal-to-noise ratio can be increased while maintaining the total noise in the system.

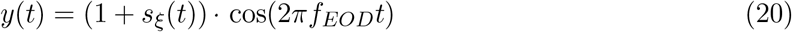

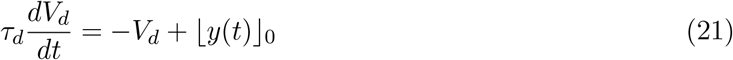

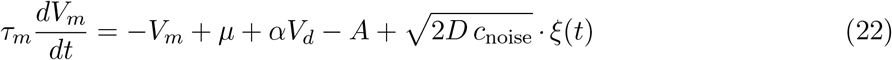

In the here used model a small portion of the original noise was assigned to the noise component (*c*_noise_ = 0.1, Fig. 10 C_iii_) and a big portion used as the signal component (*c*_signal_ = 0.9, Fig. 10 A_iii_). For the noise split to be valid (Lindner, 2022) it is critical that both components add up to the initial 100 % of the total noise and the baseline properties as the firing rate and the CV of the model are maintained. This is easily achieved in a model without a carrier if the condition *c*_signal_ + *c*_noise_ = 1 is satisfied. The situation here is more complicated. After the original noise was split into a signal component with *c*_signal_, the frequencies above 300 Hz were discarded and the signal strength was reduced after the dendritic low pass filtering. To compensate for these transformations the initial signal component was multiplied with the factor *ρ*, keeping the baseline CV (only carrier) and the CV during the noise split comparable, and resulting in *s*_*ξ*_(*t*). *ρ* was found by bisecting the plane of possible factors and minimizing the error between the CV during baseline and stimulation.

### Artificial bursts in the model

For the analysis in Fig. 5 the spikes in the non-busty model with identifier 2013-01-08-aa (see table 1 for model parameters) were supplemented by burst spikes after exactly one, two or three EOD periods. A spike was not added if the refractory time to the next spike could not be maintained.

## A Appendix

## B Acknowledgments

We are thankful for the funding of the DFG Priority Programme Evolutionary Optimisation of Neuronal Processing (SPP 2205).

